# Sexually distinct multi-omic responses to progressive endurance exercise training in the rat lung—Findings from MoTrPAC

**DOI:** 10.1101/2025.04.10.647997

**Authors:** Gina M. Many, Tyler J Sagendorf, Hugh Mitchell, James A Sanford, Samuel Cohen, Ravi Misra, Igor Estevao, Ivo Díaz Ludovico, David A Gaul, Malene E Lindholm, Mereena Ushakumary, James Pino, Nicholas Musi, Jia Nie, Facundo M Fernández, Eric A Ortlund, Karyn A. Esser, Sue C Bodine, Simon Schenk, Geremy Clair, Joshua N Adkins, The MoTrPAC Study Group

**Affiliations:** Biological Sciences Division, Pacific Northwest National Laboratory, Richland, WA USA; Division of Pulmonary and Critical Care Medicine, Cedars-Sinai Medical Center, Los Angeles, CA USA; Department of Pediatrics-Neonatology, University of Rochester-Golisano Children’s Hospital, Rochester, NY USA; School of Chemistry and Biochemistry, Georgia Institute of Technology, Atlanta, GA USA; Division of Cardiovascular Medicine, Department of Medicine, Stanford University, Stanford, CA USA; Diabetes and Aging Center, Cedars-Sinai Medical Center, Los Angeles, CA USA; Department of Biochemistry, Emory University School of Medicine, Atlanta, GA USA; Department of Physiology and Aging, University of Florida, Gainesville, FL, USA; Aging and Metabolism Research Program, Oklahoma Medical Research Foundation, Oklahoma City, OK USA; Department of Internal Medicine, Carver College of Medicine, University of Iowa, Iowa City, IA USA; Department of Orthopaedic Surgery, School of Medicine, University of California San Diego, La Jolla, CA USA; Oregon Health & Science University, Department of Biomedical Engineering, Portland, OR 97239, USA

## Abstract

Despite the lungs being essential for ventilation and aerobic exercise capacity, conventionally the lungs are not thought to adapt to exercise training. Endurance exercise is key to pulmonary rehabilitation programs, which also displays sex-specific differences in therapeutic efficacy. Given the molecular underpinnings of sex-specific lung adaptations to endurance exercise are uncharacterized, we used a multi-omics approach to study sex differences in the lungs of 6-month-old Fischer 344 rats in response to an 8 week progressive endurance treadmill training protocol. This was accomplished by reannotating publicly accessible data from the Molecular Transducers of Physical Activity Consortium (MoTrPAC) and integrating newly-analyzed acetylome data to assess multi-omic sex differences in sedentary and progressively trained states. Female rats displayed enrichment in immune-related features and pathways at the transcriptome and proteome level that were maintained with training. Conversely, in the male rat lung there was an overall decrease in immune pathways following 8 weeks of training. Sexually conserved responses to training included increased enrichment in transcriptomic pathways related to type I alveoli and proteomic pathways related to cilia, and decreased mitochondrial protein acetylation. In both sexes, features known to be enriched in lung diseases were attenuated with training. Together our findings provide novel insight into sex specific responses to endurance exercise training in the rat lung and may offer translational insight into sex-specific differences in lung disease pathogenesis and treatment.

## Introduction

While the lung is a critical conduit for ventilation and hence aerobic functional capacity during exercise, the lung is not typically thought to adapt to endurance exercise training (1,2). That said, endurance exercise training is paramount to pulmonary rehabilitation and the restoration of cardiopulmonary functional capacity following lung injury, alluding to its plasticity (3,4). Application of omics technologies have facilitated the identification of lung abnormalities associated with illnesses such as asthma, chronic obstructive pulmonary disease, and idiopathic pulmonary fibrosis (4–6). The LungMAP initiative has been pivotal in advancing our understanding of lung cellular populations. Recent progress leveraging technologies such as single-cell RNA sequencing (scRNA-seq) (7,8), sorted-cell proteomics (9), spatially resolved proteomics (10) has enabled detailed characterization of distinct cellular types within the lung, elucidating further their specific markers, functions, developmental lineages, and heterogeneity. Integration of these datasets enable identification of protein-level cell-specific markers usable for the identification of cell population shifts in proteomics datasets but also to perform immunofluorescence-based imaging using highly multiplexed protein imaging (11). Such technologies can also be applied to study the impact of external stimuli on the cellular and molecular landscape of the lung in health and disease to expand our understanding of lung physiology. To our knowledge, no studies have investigated how endurance exercise training affects the molecular landscape of the lung, especially in non-disease states–doing so offers the potential to characterize the molecular mechanisms of therapeutic lung remodeling in health and disease.

The Molecular Transducers of Physical Activity Consortium (MoTrPAC) was established to uncover molecular regulators that underpin the adaptive response to exercise, with the goal being to better understand the mechanisms by which physical activity promotes health (12). In its initial set of published studies, MoTrPAC demonstrated that exercise modulates the expression and regulation of thousands of molecules in multiple tissues, including the lung (13), and that the molecular landscape and response to training is highly tissue and sex-specific (13–16). In the lung, training was broadly found to modulate the expression of genes related to inflammation, yet interrogation of sexual dimorphism in the molecular landscape of the lung was not performed. Here we re-analyzed the lung omics datasets from the MoTrPAC data repository, expanded gene set annotation and added acetylomics profiling from the lung of these rats to provide an in-depth overview of sex-specific differences in the multi-omic landscape of the rat lung in sedentary and progressively endurance-trained states. Detailed interrogation of this dataset is intended to unveil sexual differences in the molecular landscape of the lung and identify candidate mechanisms by which exercise training may attenuate pulmonary disease risk.

## Results

### Sexual dimorphism in multi-omic landscape of the lung in sedentary rats

A study overview is provided in Fig 1A; briefly, male and female rats displayed similar cardiorespiratory improvements with training, indicated by a +14% and +17% increase in VO_2_max relative to total body mass in females and males, respectively (16). This was in contrast to −6% and −9% mL/kg/min decrease in VO_2_max in sedentary male and female rats, respectfully, over the 2 mo study protocol.

**Figure 1.**
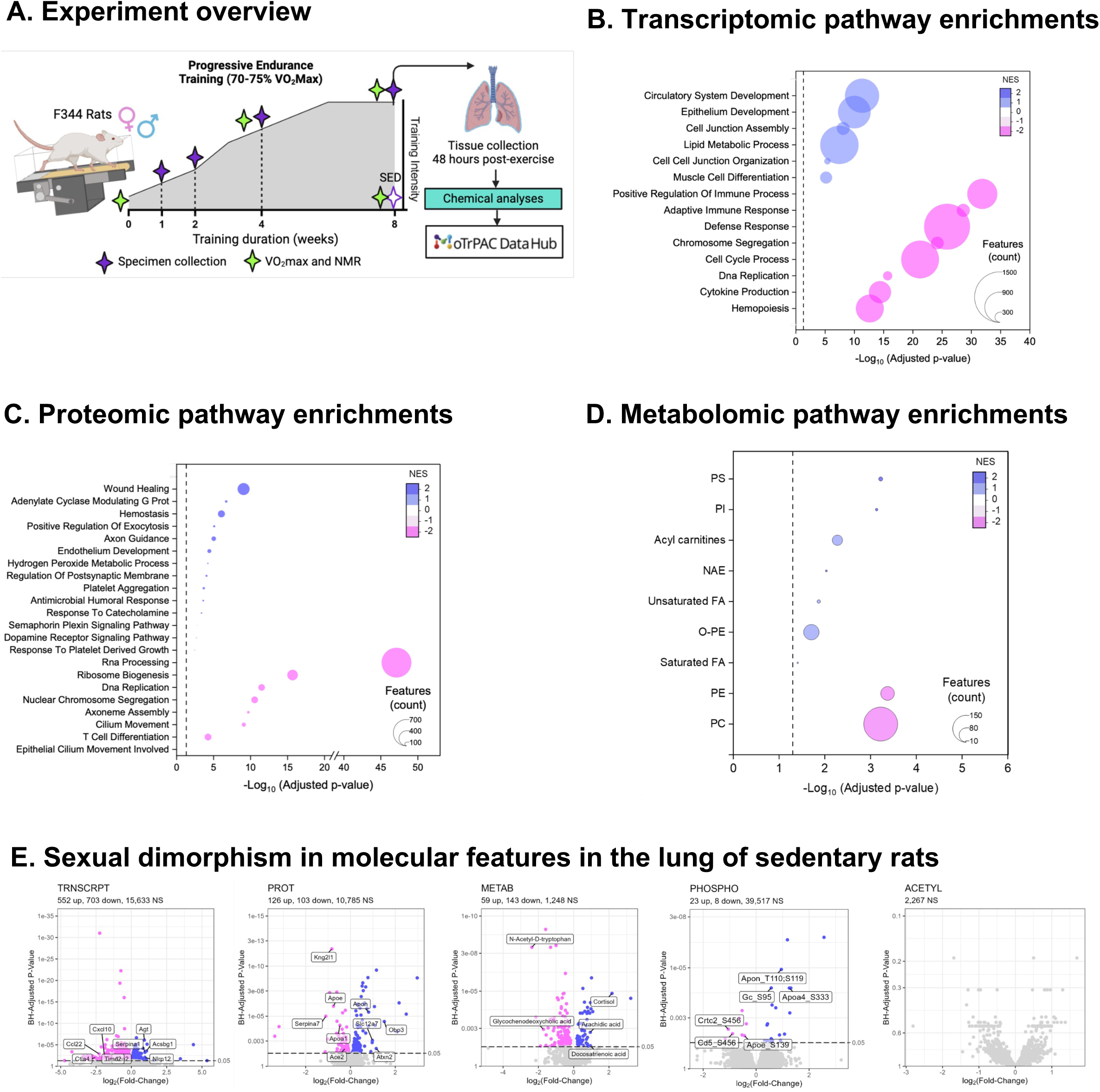
Sex differences in the multiomic landscape of the lung in sedentary rats. (**A**) Experimental overview (**B–C**) Fast gene set enrichment analysis (FGSEA) of transcriptomic (**B**) and proteomic (**C**) pathways between male and female sedentary rats, where a negative z-score indicates enrichment in females from male – female contrasts. (**D**) FGSEA of RefMet metabolite subclasses between sexes. (**E**) Volcano plots displaying features significantly enriched in the male (blue) and female (pink) sedentary rat lung at FDR <0.05.

To explore sexual dimorphism in the rat lung, we first examined sex differences in the molecular landscape of sedentary rats by re-analyzing data from the MoTrPAC data repository. To support sex-specific statistical analysis feature annotations were curated to an updated version of the rat genome (mRatBN7.2)(17). Utilizing this data, we compared the differential abundance of molecules at the transcript, protein, protein post-translational modification (PTM, phosphorylation and acetylation), and metabolite levels in the lungs of 8 month-old sedentary male and female rats. Notable sex differences were detected across all omes, with the exception of the acetylome (Table S1A-E; Fig. 1E). Specifically, 1,255 transcripts (7.4% of the measured transcriptome; 552 up, 703 down), 229 proteins (2.1%) and 202 metabolites (13.9%) were significantly different between males and females (FDR <0.05). Protein post-translational modifications showed few differences, with only 31 unique phosphosites being different between sexes (Fig. 1E).

At the transcript level, excluding Y chromosome-specific transcripts, genes with elevated expression in the sedentary male lung included *Acsbg1* (z-score: 4.8), *Serpina1 (*z-score: 4.6), *Nlrp12 (*z-score: 3.7) and angiotensinogen (*Agt,* z-score: 6.2). Whereas female-biased transcripts included the proliferation marker *Mki67* (z-score: –6.5), E-selectin (*Sele*, z-score: –6.2), *Jaml* (z-score: –5.7), and *Ccr5* (z-score: –5.3) (Fig. 1E; Table S1A). We also performed gene set enrichment analysis in the lung of sedentary rats as another method to compare sexual dimorphism in the molecular landscape of the sedentary rat lung. The female rat lung broadly displayed enrichment in transcriptomic pathways related to developmental processes and the immune system, to include terms such as adaptive immune responses, cytokine production, cell cycle processes and chromosome segregation. Males displayed enrichment in transcriptomic pathways related to circulatory and epithelial cell development, cell junctions and lipid metabolism (Fig. 1B; Table S2A).

Sex differences at the proteome level included increased abundance of endothelial cell adhesion molecule (Esam, z-score: 5.5) and von Willebrand factor A domain containing 1 (Vwa1, z-score: 4.8) in males, whereas females displayed increased protein abundance of kininogen 2 (Kng2l1, z-score: –8.2), Apoa1 (z-score: –5.2) and Apoe z-score: –6.1) (Fig. 1C&E; Table S1B). Relating to the renin-angiotensin system, vasodilatory Ace2 was enriched at both the transcript (z-score: –3.4) and protein (z-score: –3.7) level in sedentary females, whereas males displayed increased levels of Agt also at the protein level (protein, z-score: 4.6) (Fig. 1E). Global proteomic enrichment analysis displayed developmental processes, cilia and T cell differentiation enriched in females (Fig. 1C; Table S2B). Males displayed enrichments in terms related to catecholamine responses, axon guidance, and platelet aggregation (Fig. 1C). Features driving pathway enrichment in platelet activation in males included plasminogen activator, urokinase and tissue type (Plau and Plat), von Willebrand factor (Vwf), coagulation factors F9 and F11, cathepsin G (Ctsg) and vinculin (Vcl).

At the hormonal and metabolite level, the lungs of males displayed higher levels of cortisol (z-score= 5.3), docosatrienoic acid (z-score= 2.6) and arachidic acid (z-score= 3.5). The lungs of sedentary female rats displayed increases in glutathione and N-acetyl-D-tryptophan (Table S1E). Enrichment analysis according to RefMet subclass revealed an increase in acylcarnitine species, the anti-inflammatory lipid class NAE, and both unsaturated and saturated fatty acid species in the lungs of males relative to females (Fig. 1D; Table S2E). Whereas females showed enrichment in the plasmalogen PE and PC species (Fig. 1D&E).

### Multi-omic training responses in the rat lung

We next examined multi-omic responses in the rat lung after 1, 2, 4, and 8 weeks of progressive endurance exercise training where the rats reached a plateau in exercise intensity and volume during weeks 6-8 of training (Fig. 1A, (16)). All tissues were collected 48 hr after the last bout of exercise to mitigate acute effects and elucidate training-dependent effects on the lung. Over the 8-week training time course, 2,256 unique features were impacted at any given training timepoint in either sex. Overall, females showed a more robust multi-omic response to training, with 1,554 unique differential features at any training timepoint, compared to a total of 909 unique features across timepoints in males. At the transcriptomic level, this included 1,184 features in females and 321 features in males (Fig. 2A & C). Temporal dynamics of differentially regulated features differed by sex; females displayed the greatest number of differentially expressed features at the 2-week training timepoint (Fig. 2A). Males displayed the greatest number of differentially regulated features at the 1 week training timepoint, which was largely driven by differentially expressed metabolites (Fig. 2C).

**Figure 2.**
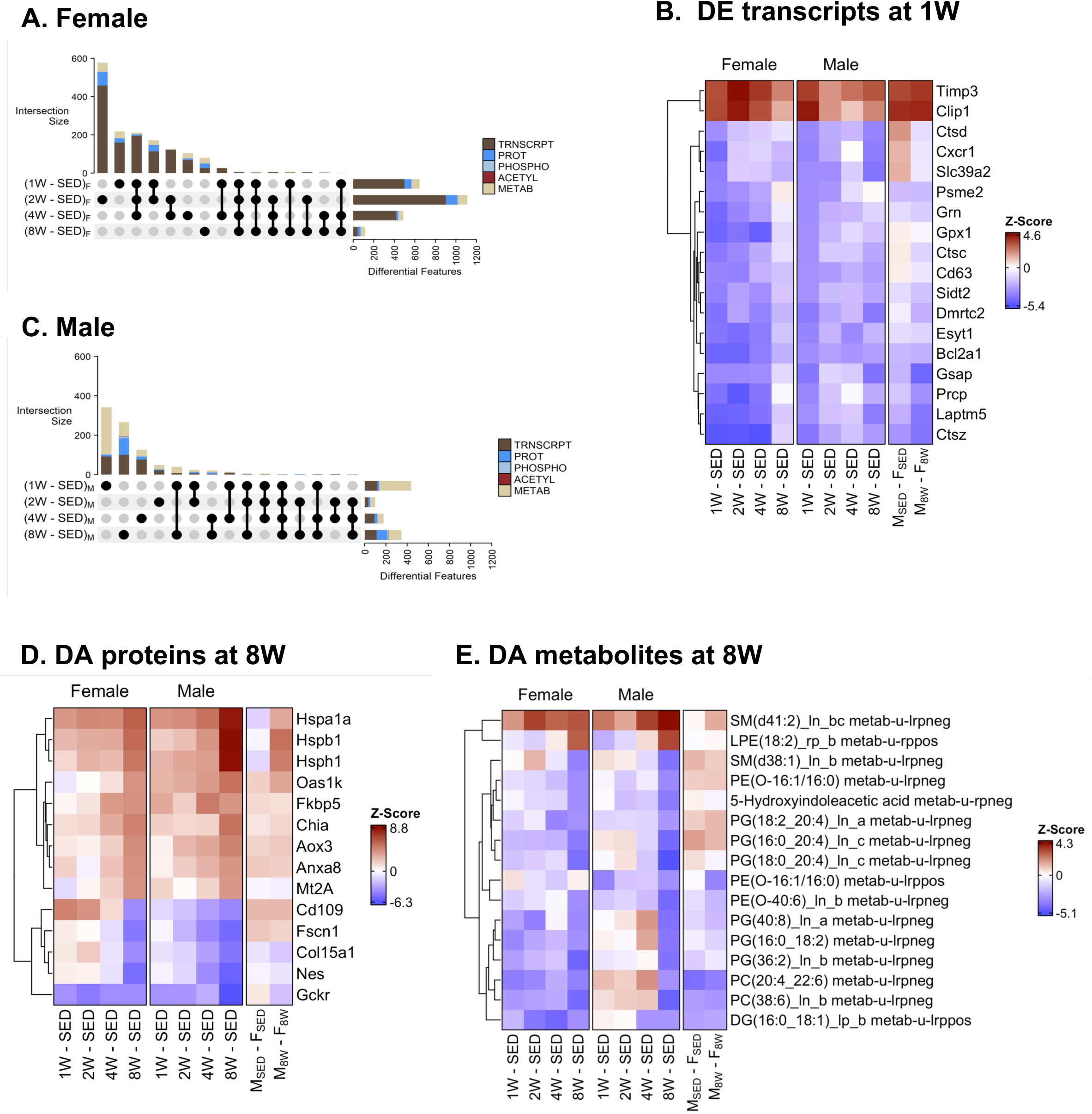
Multiomic response to progressive endurance training in the rat lung. UpSet plot displaying the number of significantly different molecular features per training time point in female (**A**) and male (**C**) rats. Here the intersection size represents the number of significantly different features per contrast, where a line connecting dots represents features shared at multiple training timepoints. (**B**) Most significant sex-consistent differentially expressed (DE) transcripts in the rat lung after 1W of training. Most significant sex-consistent differentially abundant (DA) proteins (**D**) and metabolites (**E**) after 8W of training.

*Transcriptomics.* Few sex-consistent differentially expressed genes were observed across training timepoints (Fig. 2A&C, Table S3A). In males, the greatest number of differentially expressed genes (119) were observed at the 1 week training timepoint; this time point also displayed the greatest number of shared differentially regulated genes with females. Here, both sexes displayed increased expression of *Timp3*, an inhibitor of Akt1 signaling and decreased expression of cathepsins (*Ctsc*, *Ctsd*, *Ctsz)* (Fig. 2B). At 8 weeks, males and females shared 4 differentially expressed transcripts to include an upregulation of the tumor suppressor, *Slc39a2* (Loginov et al. 2015) and *Scnn1b* involved in liquid regulation at the airway (Table S3A). Pathway enrichment analysis revealed a decrease in immune-related terms in both sexes with training, albeit the response was attenuated in 8-week-trained females, with only MHC II immune-related terms remaining decreased in 8-week-trained females. The NFkB inhibitor, *Nfkbie*, decreased throughout training in females, suggesting anti-inflammatory effects of exercise in the female lung. Both sexes displayed increased expression of transcripts involved in angiogenesis, including *Dll4*, *Efna1*, and *Efnb2* at 8 weeks in males, and *Vegfd* at 4 and 8 weeks in females. Baseline sexual dimorphism in transcriptomic pathway enrichments appeared to influence training responsiveness. For example, sedentary females displayed enrichment in transcripts related to the immune system (Fig. 3B, Msed-Fsed column, see Fig 3A), which remained enriched in females following training (Fig. 3B, Table S4A). Conversely, some sexually dimorphic pathway enrichments converged with training such as terms related to microtubule cytoskeleton organization, which were elevated in sedentary females and increased in males with training, diminishing sex-specific differences after training. With training, males but not females displayed pathway enrichment related to neuron-to-neuron synapses, which included increased expression in post-synaptic *Tamalin*, *Rapgef4*, and Inhibitory synaptic factor 1 (*Insyn1*).

**Figure 3.**
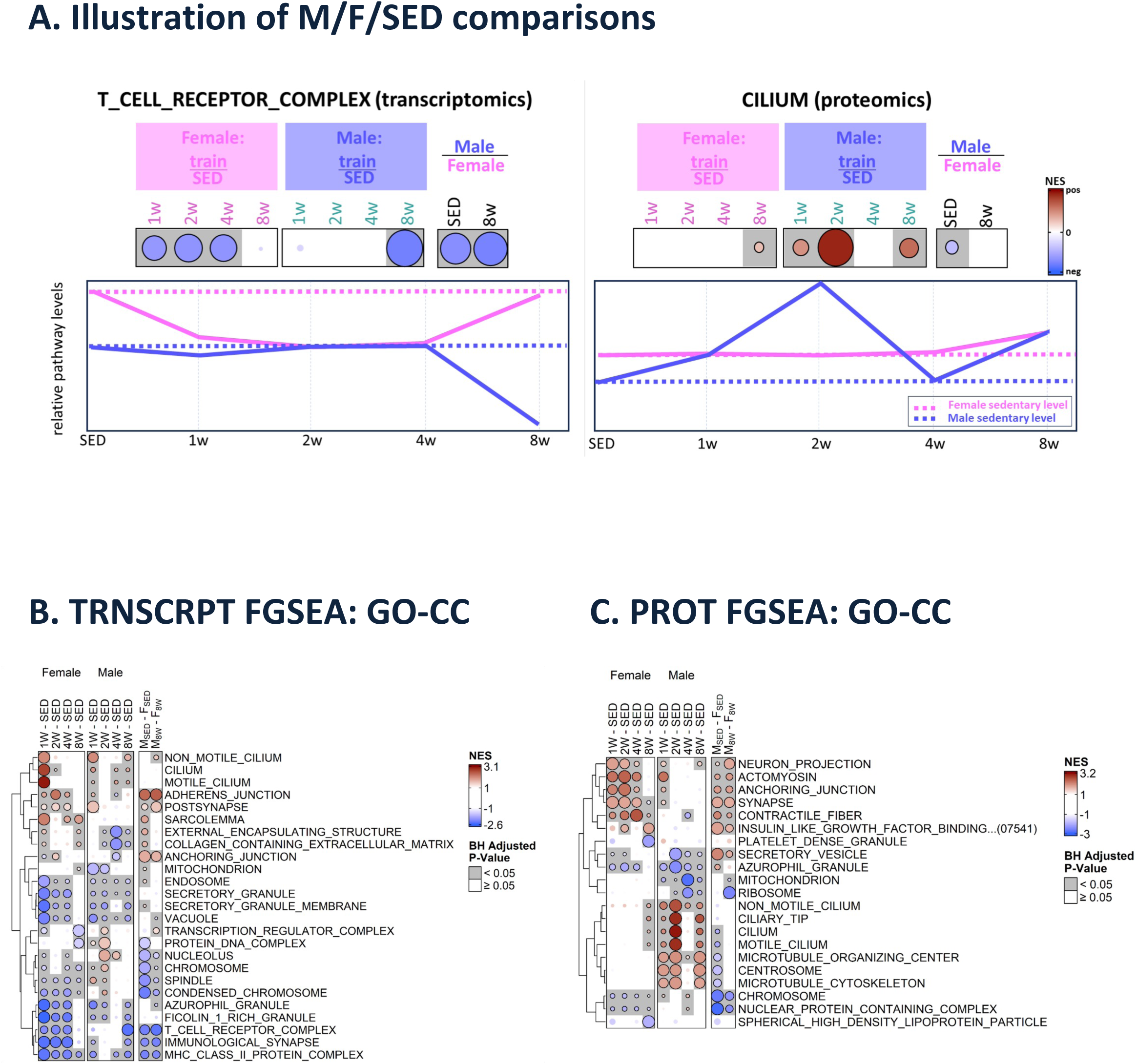
Gene set enrichment analysis across training in the rat lung. (**A**) Schematic displaying overall pathway enrichments between sedentary and trained rats, intended to clarify and illustrate the utility of the rightmost two columns of panels **B** and **C**, and other subsequent figures. Fast gene set enrichment analysis was performed by sex across training time points. Gene Ontology Cellular Component (GO-CC) enrichment results using fast gene set enrichment (FGSEA) are displayed for the (**B**) transcriptome, (**C**) proteome to include male versus female comparisons in sedentary versus 8-week-trained rat lung.

### Proteomics

Males and females shared 14 proteins with differential regulation at 8 weeks, which included enrichment of the heat shock proteins (Fig. 2D; Table S3B). In both sexes Hspa1a was upregulated at all training timepoints, while Fkbp5, an Hsp90 co-chaperone that attenuates glucocorticoid responses, was upregulated at 4 and 8 weeks in both sexes (Fig. 2D).

Interestingly, Fkbp5 is decreased in human lung cancer (18), suggesting a protective role of Hsps in lung carcinoma. Additional proteomic features that displayed sex-consistent responses to training at 8 weeks included Cd109, Col15a1, and Mt2A (Fig. 2D). Cd109, which decreased, has been implicated in adenocarcinoma of the lung (19). Col15a1, which also decreased, is expressed in vessels adjacent to major airways and is enriched in pulmonary fibrosis (20). The glucocorticoid responsive metallothionein, Mt2a, was upregulated; its downregulation has been suggested as associated with idiopathic pulmonary fibrosis (21). Together this highlights sexually conserved changes in protein abundance associated with potential protection against lung diseases.

Pathway enrichment analysis revealed the impact of exercise on shared omic responses as well as changes between sexes with training. Both sexes displayed an increase in cilia terms at 8 weeks, although the response in males began to occur at 1 week and was more robust than that of females, resulting in attenuated sex differences in ciliary terms following 8 weeks of training (Fig. 3C; Table S4B). Proteins driving enrichment included Rab23, involved in ciliogenesis (22), Dnaaf –2, –3 and –5, Ift22 and Kif3a. Exercise is known to acutely regulate cilia beat frequency (23); these results suggest that chronic exercise can affect lung ciliogenesis itself.

### Cell type enrichment analyses

Given a predominance of terms related to cellular components in global enrichment analyses, we performed cell type enrichment analysis using the HuBMAP lung single cell profiling database (24) on global transcriptomic and proteomic data to gain insight into changes in cellular compositions with training. Here, several sex-consistent patterns were observed throughout training. At the transcriptomic levels, terms related to type I pneumocytes, or alveoli, the predominant alveolar cell responsible for gas exchange in the lung increased at 1 week of training and largely remained elevated following 8 weeks of training in both sexes (Fig. 4A; Table S3A). Both male and female rats displayed a 1 week increase in terms related to submucosal gland and goblet cells that remained elevated only in males at the 8-week-trained timepoint. Overall both sexes displayed a decrease in immune cell subsets, including, B and CD4 and CD8 T cells, NK cells, alveolar macrophages, dendritic and non-classical monocytes that were significant at all training timepoints, except for 4-week-trained males (Fig. 4A).

**Figure 4.**
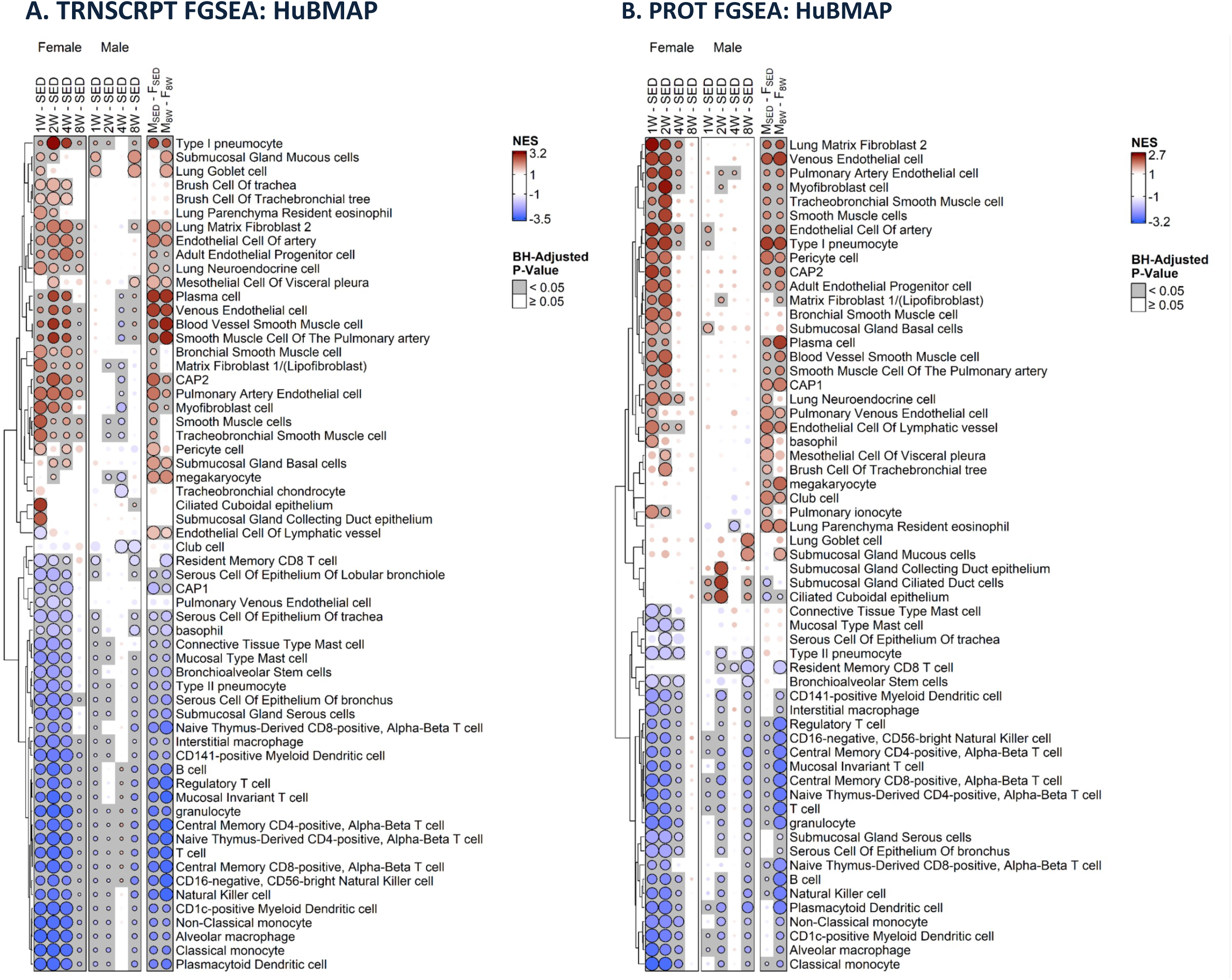
Lung cellular population gene set enrichment response to training. Fast gene set enrichment analysis was performed by sex across training time points based on HuBMAP cellular enrichments for the (**A**) transcriptome, (**B**) proteome. Scale represents normalized enrichment score (NES) where red=increased and blue=decreased gene set enrichment scaled according to intensity.

Similar patterns of sexual dimorphism and responses to exercise in cellular populations were predicted by proteomics, with sedentary males displaying higher levels of endothelial progenitors, type I pneumocytes, club cells, smooth muscle cell, pericytes and lung neuroendocrine cells that remained enriched relative to females after 8 weeks of training (Fig. 4B). Females displayed higher levels of ciliated cuboidal epithelium that were increased in males with training, resulting in attenuated differences in the 8-week-trained group. Proteomics also predicted higher abundances of leukocytes including T and B cells, granulocytes and NK cells in sedentary females that remained following 8 weeks of training.

### Metabolomic adaptations

Overall, a greater number of metabolomic terms changed with training in the male versus female lung, with males displaying 436 training-responsive features and females displaying 197 (Fig. 2A,C, Fig S1A&B). We performed enrichment analysis according to RefMet chemical subclass to assess changes in metabolite species (Fig. 5A). Shared male/female metabolite enrichments were driven by differential 15 triacylglycerol (TG) species, which were increased in males and decreased in females (Table S4E). This TG pattern corresponded to up-regulated enrichment of acylcarnitine species at 2 weeks in males, suggesting mitochondrial transport and catabolic energy conversion of excess triglycerides from 1 week (Fig. S2A). TG accumulation in the male lung attenuated at later training timepoints, coincidingly, acylcarnitine species attenuated at 4 weeks, where there was an accumulation of free carnitine (Fig. S2A, Fig. 5A). Beta oxidation proteins did not change throughout training, in fact multiple beta oxidation proteins decreased with 8 weeks of training in males rats (Fig. S2B). Conversely, cholesterol esters (CE), a minor component of lung surfactant, (Sallese et al. 2017), increased in females with training resulting in differences between sexes at 8 weeks (Table S3E & S4E, Fig. 5A). We then compared changes in surfactant-associated lipid abundance according to lipid class. Consistent with enrichment analysis, females displayed higher levels of PC, CE and cholesterol species (Fig. 5B). Given surfactant is composed of ∼10% neutral lipids this finding could reflect sex differences in neutral lipid surfactant composition or sex difference in lung-resident cellular metabolism (Sallese et al. 2017). No differences in pulmonary surfactant proteins were observed in response to training (Fig. 5C). By the 8 week training timepoint, males and females shared differential expression of 16 metabolites. Six of these species were phosphatidylglycerol (PG) lipids, which decreased (Table S3E); 5-hydroxyindoleacetic acid, the primary metabolite of serotonin breakdown, decreased in the lung in both male and female rats at 8 weeks (Fig. 1E; Table S3E). Interestingly, reduced clearance of serotonin is associated with accelerated lung injury (25), further highlighting effects of exercise on the lung that are inversely associated with disease risk.

**Figure 5.**
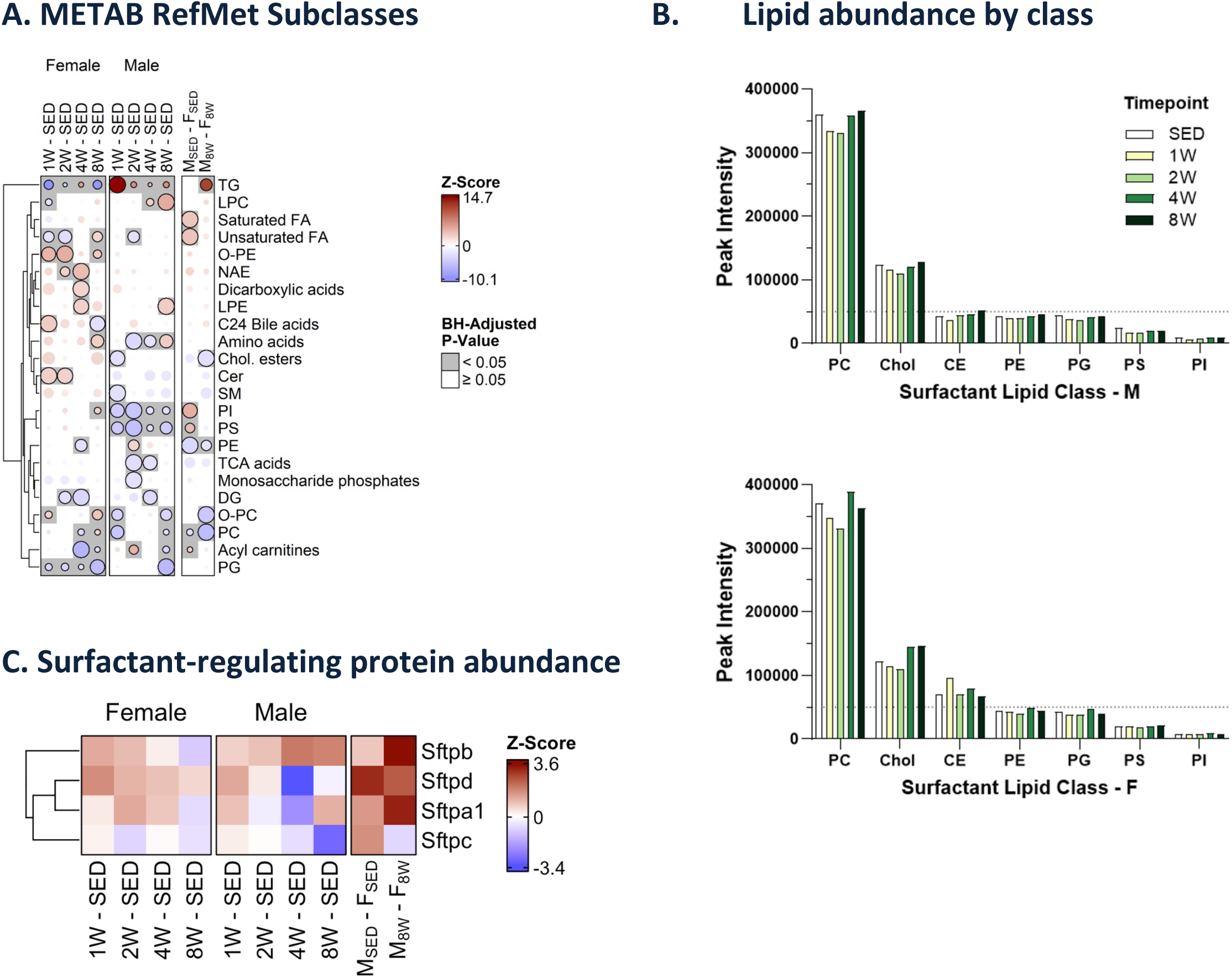
Metabolomic and lipidomic responses to training in the rat lung. (**A**) Gene set enrichment analysis according to RefMet subclass. (**B**) Total abundance of surfactant-associated lipid subclasses based on normalization to internal standards during LC-MS/MS. (**C**) Changes in surfactant-related proteins.

### Changes in protein post-translational modifications

We utilized an integrative multi-PTM platform to further examine the multi-omic response to exercise training in the lung and inform differential signaling regulation.

*Acetylation*. With training, increased Hspa1a protein acetylation in 8-week-trained males was the only detectable change with training after correction for multiple hypothesis testing (Fig. S1B). Pathway enrichment analysis revealed decreased acetylation of mitochondrial-related proteins with 8 weeks of training in both sexes (Fig. 6A; Table S4C). Acetylsites driving enrichment in males included Hadha K214, K436 and K406, Suclg1 K57, Dbt K304 and Acat1 K400 and K187. In females driving acetylsites included Cs K316, Idh3b K374, Hadha K436, K202 and K192, At5po K100 and Suclg1 K66. In the skeletal muscle, mitochondrial protein acetylation appears related to functional capacity, as mice with high running capacity have decreased acetylation of mitochondrial proteins that is further decreased with exercise (26); this suggests mitochondrial protein acetylation may also increase metabolic efficiency in the lung.

**Figure 6.**
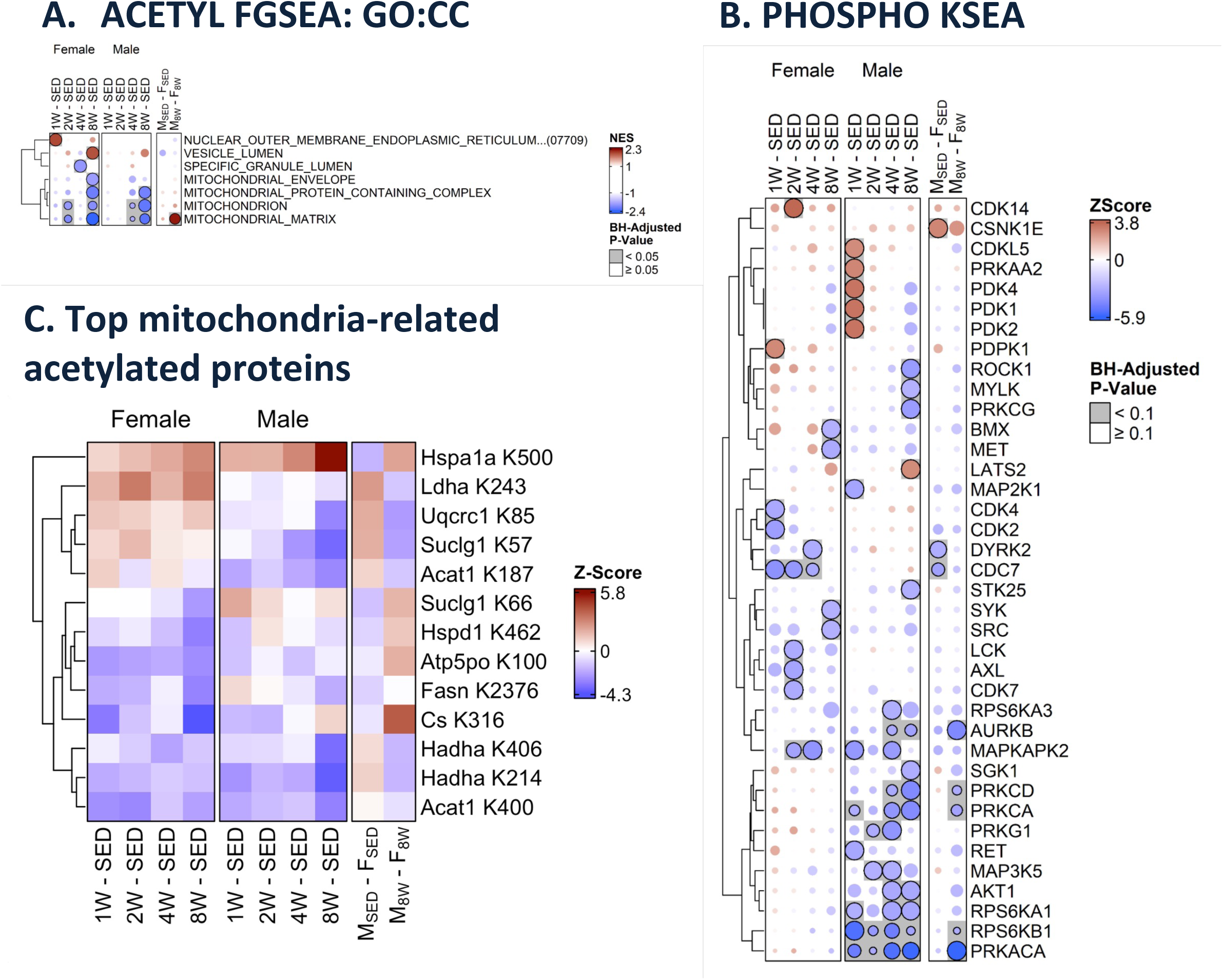
Changes in protein post-translational modifications in the rat lung with training. (**A**) FGSEA of acetylomic data. (**B**) Kinase substrate enrichment analysis (KSEA) of phosphoproteomic data curated according to PhosphositePlus. (**C-D**) Changes in top training differentially regulated mitochondrial acetylated proteins.

*Phosphorylation*. Few phosphoproteins were differentially expressed in the lung with training. Despite few phosphosites changing passing our significance threshold, we examined predicted changes in in kinase activity through kinase substrate enrichment analysis (Fig. 6B) (27). We first examined the impact of sex on predicted kinase activity in the lungs of sedentary rats. We observed predicted enrichment in CDC7 and DYRK2 in female rats, both of which decreased with training in females but not males resulting in comparable predicted activity following 8 weeks of training (Fig. 6B, Table S4D). In male rats, CSNK1E activity was elevated in sedentary rats but such differences were attenuated with training. Both male and female rats displayed decreases in MAPKAPK2 during intermediate training timepoints. Interestingly, we also observed down-regulation of PRKACA (protein kinase A or PKA) and AKT1 in males at multiple training timepoints, both of which have been shown to antagonize ciliogenesis (Mao et al. 2019; Senatore et al. 2021). Since we observed enrichment of cilium-related terms in up-regulated proteins, this is consistent with a reciprocal relationship between these signaling moderators, and cilium development during long-term exercise training.

## Discussion

Despite the lung being central to cardiorespiratory fitness and that endurance exercise is a key component of pulmonary rehabilitation, little knowledge of the sex-specific impact of endurance exercise on the molecular landscape of the lung exists, especially in non-diseased lungs. To this end, MoTrPAC was designed to uncover the molecular regulators of adaptive responses to exercise, and by extension, establish the mechanisms by which physical activity promotes health and combats disease. As emphasized by increased severity of disease outcomes in males during the SARS-CoV2 pandemic, sex differences in the molecular landscape of tissues, including the lung, likely drive differences in disease outcomes and risk (28). We thus leveraged the MoTrPAC data repository to examine sex differences in the molecular landscape of the lung as well as its temporal response to progressive endurance exercise training. Our findings highlight key sex differences in the molecular landscape of the lung, indicative of enrichment of pathways related to a variety of immune cell subsets in the lungs of female rats and enrichment in features relating to airway structural components in the sedentary male rat lung. With training, both male and female rats displayed a decrease in immune cell pathway enrichments, albeit more robust in males, suggestive of anti-inflammatory effects of exercise on the lung. Sex consistent responses to 8 weeks of endurance exercise training included pathway enrichments relating to ciliogenesis and alveolar type I cells, highlighting the ability of prolonged endurance exercise to beneficially structural components of the lung. Further, endurance training shifted the abundance several genes and proteins to inversely correlate to disease. Together, our findings highlight novel mechanisms by which endurance exercise remodels the lung to protect against disease.

The respiratory system of males and females differs in terms of development and disease susceptibility. In both humans and rats, females displays accelerated lung development *in utero* largely attributed to the roles of estrogen (29,30). Postnatally, sex differences are also observed. In humans for example, males have larger lungs and conducting airways compared to females, where female lungs have more abundant yet smaller airways (31,32). Rodent studies suggest that estrogen contributes to structural differences in the lung as ovariectomy attenuates reduced alveolar size that is observed in females (33). The risk and outcomes of pulmonary diseases also differs by sex. Female smokers, for example, have increased risk of smoking-induced airway constriction and COPD relative to males even at lower exposure levels (34,35). Rodent studies also show increased susceptibility to cigarette smoke exposure in females (36), where female mice show increased small airway remodeling and oxidative stress that is alleviated by ovariectomy (37). Females additionally have a higher risk of asthma and associated disease severity (38). Contrastingly, males display increased emphysema severity (39). It is thus likely that sex hormones, and potentially other factors like genetic imprinting and embryonic cellular establishment, impact respiratory disease susceptibility and outcomes, with females appearing overall more susceptible to reactive airway constriction. Other notable sex differences in the multiomic landscape of the lung include males-specific enrichments in the renin-angiotensin system. Males displayed higher levels of vasopressive Ace and Agt, whereas females displayed increases in vasodilatory Ace2. The risk of hypertension is greater in males compared to premenopausal females, an effect suggested to be driven by estrogen. However, during the menopausal transition such differences in the incidence of hypertension are attenuated–yet the response to anti-hypertensive treatment differs between males and females, with females displaying overall reduced treatment efficacy and increased side effects. Despite this knowledge, the molecular and cellular regulators of sexual dimorphism in the lung are relatively unexplored and can provide critical information regarding sex-specific precision medicine approaches for lung disease treatment.

While the molecular mechanisms driving sex differences in disease risk and outcomes are largely unknown, in the current study we identify that differences in lung resident immune cellular populations that provide insight into differences in disease pathogenesis. We observed the female rat lung displayed enrichment in molecular features predictive of immune cell populations to include T and B cells, NK cells, monocytes and dendritic cells in both sedentary and progressively trained states. Whereas, sedentary males displayed enrichment in platelet activation and megakaryocyte terms. In the lung, megakaryocytes, the cellular precursor of platelets, are shown to have distinct inflammatory profiles (40). Both megakaryocyte and platelets can be infected with SARS-CoV2, resulting in increased reactivity (41). Male-specific enrichments in lung megakaryocyte populations could contribute to worsening of SARS-CoV2 outcomes in males; a hypothesis warranting future investigation (28). In females, our findings of predicted increases in immune cell populations also pair with sex differences in disease risk. Females display elevated autoimmune disease risk, including those affecting the lung such as lupus, autoimmunity-associated shrinking lung syndrome and idiopathic pulmonary fibrosis (38), yet beneficially display a more robust immune response to lung cancer (35,42). Such differences in disease risk and response could be attributed to an increased abundance of resident immune cell populations in the female lung.

Sex dimorphism in immune-related responses to endurance training was also observed, with males displaying more robust anti-inflammatory effects of endurance training evident by reductions in multiple immune pathways in the 8-week-trained male rat lung, with females showing decreases primarily in immune pathways related to MHC II protein complex following 8 weeks of training. Sex conserved responses suggestive of MHC II protein complex suppression following 8 weeks of training is suggestive of attenuated immunoreactivity. This training response could be linked to observed enrichments in ciliary features following 8 weeks of training in both sexes, as increased mucus clearance could reduce overall exposure to respiratory pathogens. While moderate levels and intensities of endurance training are attenuate acute respiratory illness risk (43), the benefits of exercise on mucosal clearance are thought to be driven by increased airway shear stress (44). Our findings suggest that regular endurance training may protect the lung from pathogens and particulates by directly regulating ciliogenesis, thereby improving lung mucociliary clearance and thereby reducing reactivity of resident immune cells. A candidate mechanism by which exercise promotes ciliogenesis includes Timp3-mediated inhibition of Akt1 signaling. This is supported as *Timp3* expression increased in both male and females rodents throughout training and *Timp3* inhibits Akt1 signaling, an antagonist of ciliogenesis (45,46). Phosphoproteomics data predicted a decrease in Akt1 signaling with training, which was only observed in males; lack of an effect in females may be driven by immune cell enrichments in Akt1.

### Summary

Our findings suggest that endurance training leads to beneficial effects in the lung of both sexes, as indicated by a shift in molecular signatures linked to lung disease pathogenesis and beneficial structural remodeling to include ciliogenesis. Our work adds to a body of literature supporting the need to stratify for sex when studying responsiveness to interventions and therapies and that sex differences in the molecular landscape of the lung exist, which have future implications for targeted treatment approaches.

## Software

The R programming language was used for all statistical analyses and data visualization (R Core Team 2024). The following R/Bioconductor (Huber et al. 2015) software packages were used extensively: Biobase (v2.66.0) (Huber et al. 2015), ComplexHeatmap (v2.22.0) (Gu 2022), edgeR (v4.4.0) (Chen et al. 2024), fgsea (v1.32.0) (Korotkevich et al. 2016), limma (v3.62.0) (Ritchie et al. 2015), tidyverse (v2.0.0) (Wickham et al. 2019) and TMSig (v1.0.0) (Sagendorf et al. 2024).

## Methods

### Experimental Design

Full details of the design of this MoTrPAC study have been published previously (13,14,16) and are available via the MoTrPAC website www.motrpac.org). Briefly, 6-month male and female Fischer 344 inbred rats obtained from the National Institute on Aging (NIA) rodent colony were familiarized to treadmills, and compliant animals were randomized to control or exercise intervention groups. Endurance exercise training consisted of 5 consecutive days per week at 70-75% VO_2_max using a progressive treadmill protocol, with animals undergoing 1, 2, 4 or 8 weeks of training. Sedentary control animals were placed on treadmills for 15 min/day at a speed of 0 m/min for 5 consecutive days per week for a total of 8 weeks. Body composition was determined for all rats 13 days prior to the start of training using the minispec LF90II Body Composition Rat and Mice Analyzer (Bruker), with post-training body composition determined for rats in the control, 4-week and 8-week trained groups 5 days prior to sacrifice. VO_2_max was determined prior to the onset of training in all rats and during the last week of training for the control, 4-week and 8-week groups. Upon training completion, tissues were collected 48 hours after the last treadmill intervention. All tissues were flash frozen in liquid nitrogen immediately upon removal and stored at –80°C prior to shipment to the MoTrPAC Biorepository.

### Multi-omic Data Generation

This manuscript leverages datasets generated by The MoTrPAC Study Group from male and female rat lung. Full details on the experimental methods used for processing samples and collecting data from RNA sequencing-based transcriptomics, LC-MS/MS-based proteomics, and multiple targeted and untargeted metabolomics and lipidomics platforms have been extensively described elsewhere (13,14). Quantitative metabolomics and lipidomics datasets studied here are the same as those used in previous work and publicly available through the MoTrPAC Data Hub. Importantly, the raw transcriptomics and proteomics data previously generated were reanalyzed for this work, utilizing the most recent mRatBN7.2 transcript and protein collections released in 2020 (17) and available from NCBI RefSeq. Raw data was processed by the MoTrPAC Bioinformatics Center using identical pipelines as previously used with the Rnor_6.0 reference sequences, and the resulting datasets were further processed using the same methods for quality control inspection, outlier identification, normalization and batch correction as previously performed.

### Differential Analysis

Differential analysis was performed with the edgeR and limma R/Bioconductor software packages as described previously (14). RNA-Seq data was prepared by removing low-count transcripts with the edgeR::filterByExpr function and calculating library size scaling factors with edgeR::normLibSizes. Since variability was observed among biological replicates, edgeR::voomLmFit was used to combine observation-level weights with sample-level quality weights to down-weight low-abundance observations and all observations from more variable samples (47,48). Proteomics, phosphoproteomics, and metabolomics datasets were analyzed with limma. As with RNA-Seq, biological replicates appeared to cluster poorly, so sample-level quality weights were computed with limma::arrayWeights (method=”genebygene”) and incorporated into the linear model (49). For proteomics, phosphoproteomics, and metabolomics datasets, a no-intercept model was constructed with the experimental group as the predictor (each combination of sex and timepoint) with limma::lmFit. RNA integrity number, median 5’-3’ bias, percent of reads mapping to globin, and percent of PCR duplicates as quantified with unique molecular identifiers were included as covariates in the RNA-Seq model after they had been mean-imputed and standardized (13). After fitting the models, sex-specific trained vs. SED contrasts (e.g., F.1W – F.SED) and contrasts comparing males to females at the SED and 8W timepoints were generated with limma::contrasts.fit. Robust empirical Bayes moderation was carried out with limma::eBayes to squeeze the residual variances toward a common value (RNA-Seq) or a global trend (proteomics, phosphoproteomics, and metabolomics). The resulting p-values were adjusted across sets of related comparisons (i.e., across all male vs. female or trained vs. SED contrasts) by ome using the Benjamini and Hochberg (BH) method to control the false discovery rate (FDR). Features were considered to be sufficiently differentially expressed/abundant if the FDR was <5%.

### Enrichment Analysis

The differential analysis results tables for each combination of tissue and ome were converted to matrices of z-scores with either gene symbols, RefMet metabolite/lipid IDs, or singly-phosphorylated/acetylated peptides as rows and contrasts as columns. Conversion from original to target features was carried out by selecting the most extreme z-score for each combination of contrast and target feature. These matrices serve as input for the enrichment analyses.

Gene Ontology gene sets from the C5 collection of the Molecular Signatures Database (MSigDB; v2023.2.Hs) and a single-nuclei database (24). Metabolites and lipids were grouped according to chemical subclasses from the RefMet database, while protein phosphorylation sites were grouped according to their known human protein kinases provided in the “Kinase_Substrate_Dataset” file from PhosphoSitePlus (PSP; v6.7.1.1; https://www.phosphosite.org/staticDownloads.action; last modified 2023-11-17) (27).

For each combination of tissue and ome, molecular signatures were filtered to only those genes, metabolites/lipids, or phosphosites that appeared in the differential analysis results. After filtering, all molecular signatures were required to contain at least 5 features, with no maximum size restrictions. Then, groups of highly similar Gene Ontology gene sets were identified with the clusterSets function from the TMSig R/Bioconductor package (51). This function utilizes the same clustering procedure utilized by the MSigDB to reduce redundancy within the Gene Ontology and Reactome databases. Once the clusters were identified, the largest gene set from each cluster was retained for analysis. Ties in the maximum sizes were broken using the following sequential criteria: 1) highest overlap with the background set of genes measured in the omics data, 2) gene set descriptions containing “lung” or “respir”, 3) shortest set description (i.e., broader terms), and 4) first according to alphabetic order of descriptions.

Analysis of the molecular signatures was carried out with fast gene set enrichment analysis (FGSEA) using the fgsea R package using the TMSig R/Bioconductor package (51). P-values were adjusted across all male vs. female or trained vs. SED contrasts separately by database using the method of Benjamini and Hochberg to control the FDR. The *enrichmap* function from the TMSig R package was used to generate bubble heatmaps of the ∼60 terms with the highest median –log10 adjusted p-value across contrasts (51) and using curated terms from the results.

## Supplementary Materials

### Supplementary Figures

**Figure S1.**
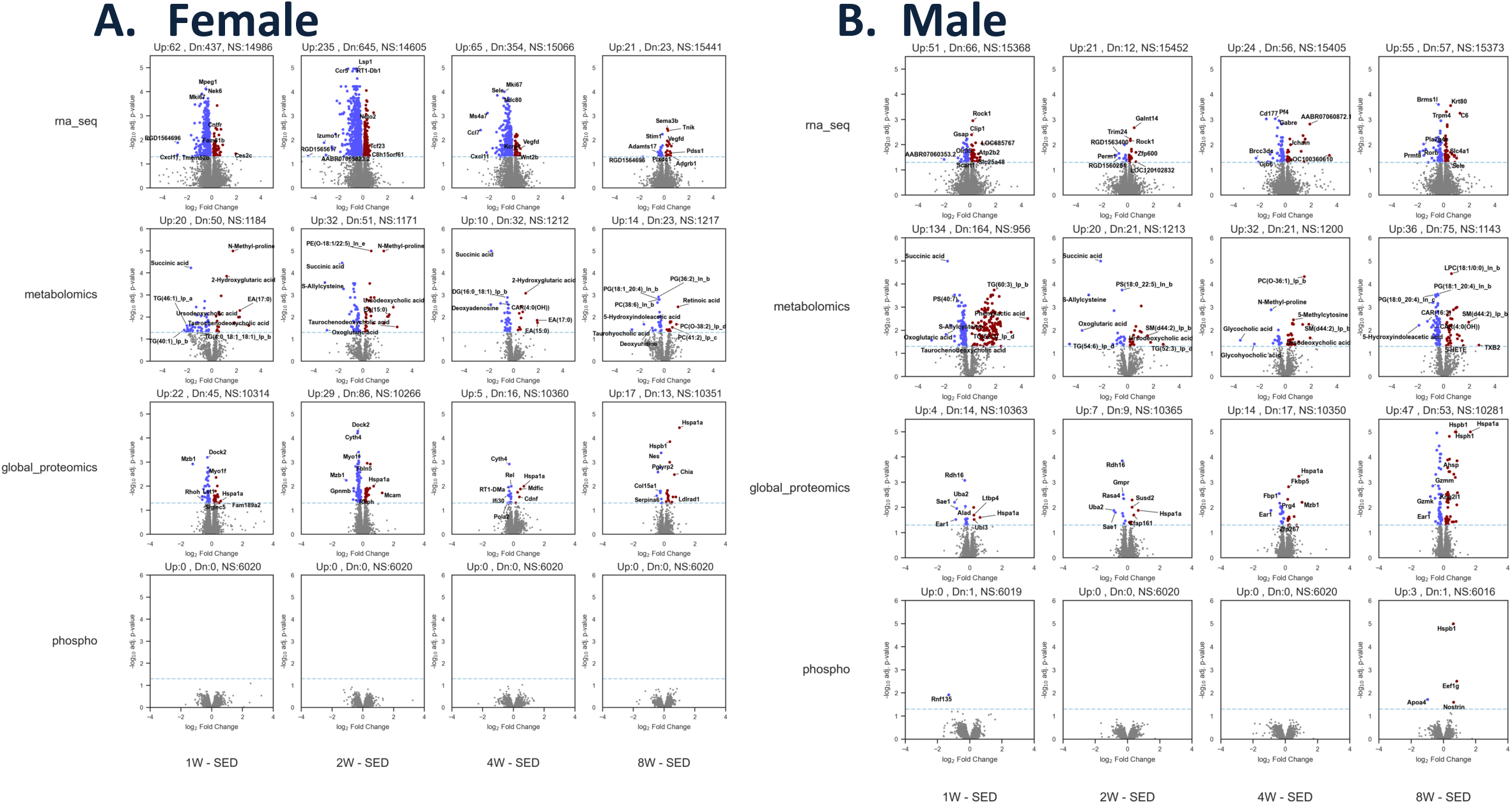
Volcano plots displaying differential multi-omic features induced in response to 1, 2, 4 or 8 weeks of endurance exercise training in female. (**A**) and male (**B**) rats.

**Figure S1.**
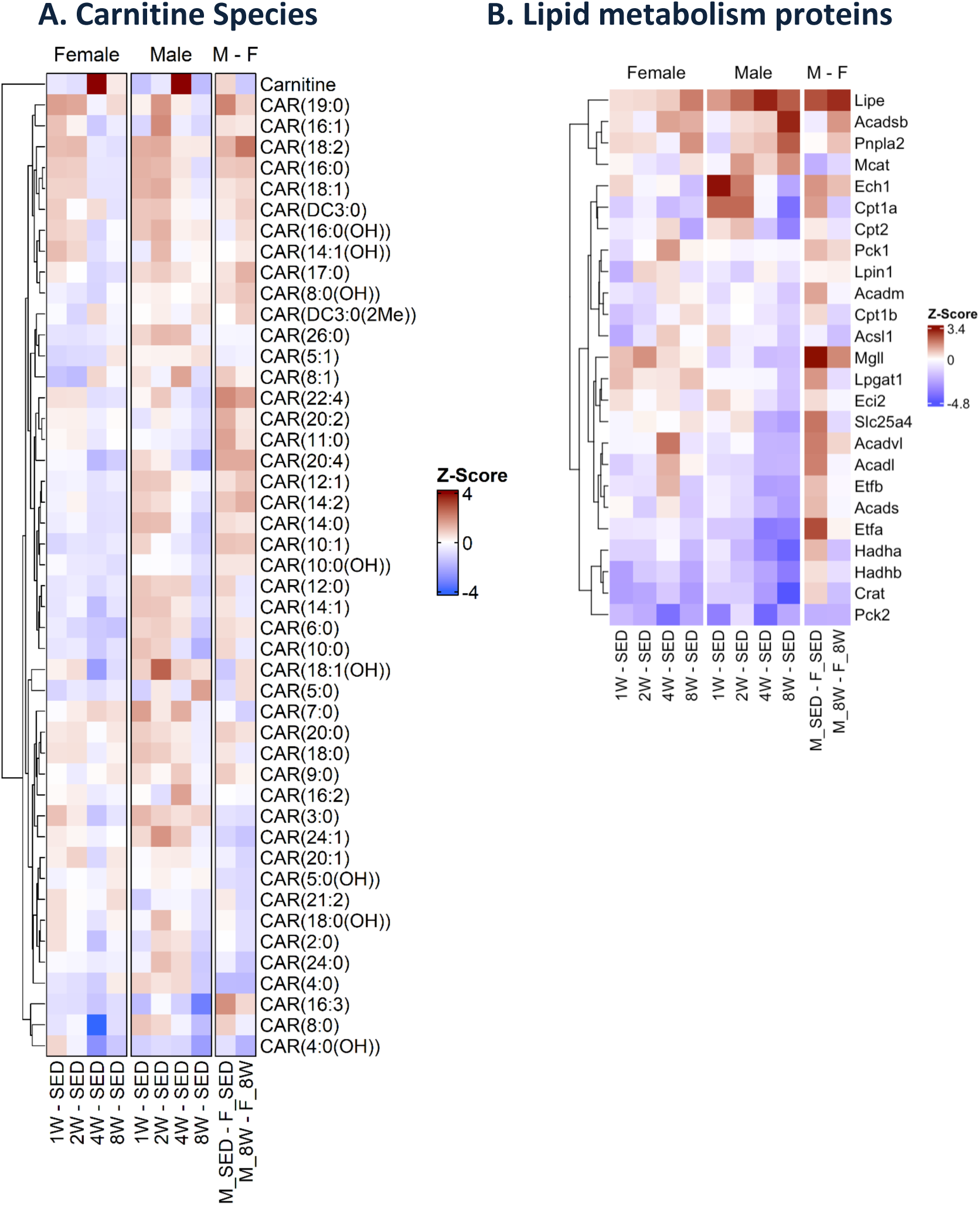
Heatmaps displaying. (A) carnitine and acycarnitine species, (B) lipid metabolism proteins.

## Supplementary Tables

Supplemental Table: Omics DA and enrichment data

## *MoTrPAC Study Group Author List

Marina Gritsenko^1^, Chelsea Hutchinson-Bunch^1^, Vladislav A. Petyuk^1^, Paul D. Piehowski^1^, Wei-Jun Qian^1^, Michael E. Miller^13^, Bridget Lester^14^, Scott Trappe^14^, Todd A. Trappe^14^, Steven Carr^15^, Pierre M. Jean Beltran^15^, Hasmik Keshishian^15^, Charles C. Mundorff^15^, Chih-Yu Chen^16^, Zhenxin Hou^16^, Xueyun Liu^16^, Kristal M. Maner-Smith^16^, Carter Asef^17^, Samuel G. Moore^17^, Mary Anne S. Amper^18^, Yongchao Ge^18^, Kristy Guevara^18^, Minghui Lu^18^, Nada Marjanovic^18^, Venugopalan D. Nair^18^, Nora-lovette Okwara^18^, Stas Rirak^18^, Yifei Sun^18^, Mital Vasoya^18^, Alexandria Vornholt^18^, Xuechen Yu^18^, Laurie J. Goodyear^19^, Michael F. Hirshman^20^, Sarah J. Lessard^21^, Brent G. Albertson^20^, Nathan S. Makarewicz^20^, Ashley Xia^22^, Marcas M. Bamman^23^, Thomas W. Buford^24^, Gary R. Cutter^25^, Kerrie L. Moreau^26^, Matthew Babcock^26^, Bryan C. Bergman^26^, Daniel H. Bessesen^26^, Zachary Clayton^26^, Catherine M. Jankowski^26^, Wendy M. Kohrt^26^, Edward L. Melanson^27^, Irene E. Schauer^27^, Dam Bae^28^, Kyle S. Kramer^10^, Ana K. Lira^10^, Blake B. Rasmussen^29^, Cynthia L. Stowe^30^, Scott Rushing^13^, Michael P. Walkup^13^, Jimmy Zhen^31^, David Jimenez-Morales^31^, Natarajan Raja Archana^31^, Michael P. Snyder^32^

^1^Biological Sciences Division, Pacific Northwest National Laboratory, Richland, WA USA

^2^Division of Pulmonary and Critical Care Medicine, Cedars-Sinai Medical Center, Los Angeles, CA USA

^3^Department of Pediatrics-Neonatology, University of Rochester-Golisano Children’s Hospital, Rochester, NY USA

^4^School of Chemistry and Biochemistry, Georgia Institute of Technology, Atlanta, GA USA

^5^Division of Cardiovascular Medicine, Department of Medicine, Stanford University, Stanford, CA USA

^6^Diabetes and Aging Center, Cedars-Sinai Medical Center, Los Angeles, CA USA

^7^ Department of Biochemistry, Emory University School of Medicine, Atlanta, GA USA

^8^Department of Physiology and Aging, University of Florida, Gainesville, FL, USA

^9^Aging and Metabolism Research Program, Oklahoma Medical Research Foundation, Oklahoma City, OK USA

^10^Department of Internal Medicine, Carver College of Medicine, University of Iowa, Iowa City, IA USA

^11^Department of Orthopedic Surgery, School of Medicine, University of California San Diego, La Jolla, CA USA

^12^Oregon Health & Science University, Department of Biomedical Engineering, Portland, OR 97239, USA

^13^School of Medicine, Wake Forest University, Winston-Salem, NC USA

^14^Ball State University, Muncie, IN USA

^15^Broad Institute, Broad Institute of MIT & Harvard (Carr Lab), Cambridge, MA USA

^16^Emory University, Atlanta, GA USA

^17^Georgia Institute of Technology, Atlanta, GA USA

^18^Icahn School of Medicine at Mount Sinai, New York City, NY USA

^19^Joslin Diabetes Center, Harvard Medical School, Boston, MA USA

^20^Section on Integrative Physiology and Metabolism, Joslin Diabetes Center, Harvard Medical School, Boston, MA USA

^21^Fralin Biomedical Research Institute/Virginia Tech, Joslin Diabetes Center, Boston, MA USA

^22^Division of Diabetes, Endocrinology, & Metabolic Diseases, National Institute of Diabetes and Digestive and Kidney Diseases, National Institutes of Health, Bethesda, MD USA

^23^The University of Alabama at Birmingham, Birmingham, AL USA

^24^The University of Alabama at Birmingham & Birmingham/Atlanta VA GRECC, Birmingham AL USA

^25^School of Public Health, The University of Alabama at Birmingham, Birmingham, AL USA

^26^University of Colorado, Denver, CO USA

^27^Anschutz Medical Campus, University of Colorado, Denver, CA USA

^28^University of Iowa, Iowa City, IA

^29^University of Texas Health San Antonio, University of Texas Health Science Center, San Antonio, TX USA

^30^Biostatistics and Data Science, School of Medicine, Wake Forest University, Winston-Salem, NC USA

^31^Department of Medicine, Stanford University, Stanford, CA USA

^32^Department of Genetics, Stanford University, Stanford, CA USA

